# Eccentric cycling involves greater mental demand and cortical activation of the frontoparietal network

**DOI:** 10.1101/2022.11.05.515299

**Authors:** Lénaic Borot, Benjamin Pageaux, Davy Laroche, Grégoire Vergotte, Ro-muald Lepers, Stéphane Perrey

**Author notes:** **Corresponding author:** Prof. Stephane Perrey (PhD) EuroMov Digital Health in Motion 700 avenue du Pic Saint Loup 34090 Montpellier France.

## Abstract

Eccentric, compared to concentric exercise, is proposed to involve different neuro-motor processing strategies and a higher level of mental demand. This study compared eccentric and concentric cycling at matched perceived effort and torque for the mental demand and related-cortical activation patterns. Nineteen men (30 ± 6 yrs) performed four different 5-min cycling conditions at 30 RPM on a semi-recumbent isokinetic cycle ergometer: 1) concentric at a moderate perceived effort (23 on the CR100® scale) without torque feedback; 2) concentric and 3) eccentric at the same average torque produced in the first condition; and 4) eccentric at the same moderate perceived effort than the first concentric condition. The conditions 2-4 were randomised. After each condition, mental demand was monitored using the NASA Task Load Index scale. Changes in oxy-(O_2_Hb) and deoxy-(HHb) haemoglobin during exercise were measured over both prefrontal cortices and the right parietal lobe from a 15-probe layout using a continuous-wave NIRS system. Mental demand was significantly higher during eccentric compared to concentric cycling (+52%, *p* = .012) and when the exercise intensity was fixed by the torque rather than the perceived effort (+70%, *p* ˂ .001). For both torque- or perceived effort-matched exercises, O_2_Hb increased significantly (*p* < .001) in the left and right prefrontal cortices, and right parietal lobe, and HHb decreased in the left, and right, prefrontal cortices during eccentric compared to concentric cycling. This study supports that acute eccentric cycling, compared to concentric cycling, involves a higher mental demand and frontoparietal network activation.

## INTRODUCTION

Most human movements involve both eccentric and concentric muscle contractions (e.g., walking, running, jumping) and a large proportion of our daily living activities require accurate control of movements employing eccentric contractions (e.g., walking downstairs, sitting, lowering weight). During eccentric contractions, the external force on the muscle is greater than the force that the muscle can generate, producing a lengthening contraction; and vice versa for concentric contractions.^1^ In the last decades, eccentric cycling at moderate intensity has been shown to promote functional (knee-extensor strength) and structural (hypertrophy) muscle adaptations in different populations due to its capability to achieve high loads on muscle tissue at low cardiorespiratory stress (athletic^2^ and chronic diseases^3^).

Over the last 60 years, increasing research has been undertaken to understand mechanical, peripheral and neural features, as well as the metabolic cost of eccentric contractions. Eccentric cycling was initially introduced to investigate the physiology of concentric and eccentric muscle work^4^. Eccentric cycling requires substantially less oxygen uptake than concentric cycling to develop the same absolute external mechanical power output.^4,5^ In addition, corticospinal excitability, electromyographic activity and rating of perceived effort are lower during eccentric compared to concentric cycling at the same absolute mechanical workload.^5–7^ Notably, at a given metabolic workload, heart rate and rating of perceived effort were found higher in eccentric compared to concentric cycling.^5,6^

Eccentric cycling is associated with substantial effort perception and sustained attention when subjects are asked to match a given target mechanical power output.^8,9^ However, little is known about cortical activity patterns of attentional resources involved during eccentric cycling. Current literature contains several lines of evidence indicating that neurobehavioral relationships in eccentric contractions are associated with higher processing demands by various brain areas for controlling motor output.^10^

At the behavioural level, several studies have reported greater variability in the motor output during eccentric compared to concentric contractions.^11^ This increased motor variability may indicate less practice and that eccentric contractions might be more cognitively demanding than concentric contractions.^10^ This hypothesis is supported by the recent findings from Kan et al.^9^ during submaximal eccentric and concentric cycling. First, an overall higher perception of the workload, evaluated by the NASA task load index (NASA-TLX) questionnaire, was reported during eccentric compared to concentric cycling. This difference is specifically related to two items of the questionnaire: mental demand and frustration. Second, the average dual-task reaction time in assessing cognitive load was 15% slower, and the accuracy score was 7% lower, during eccentric compared to concentric cycling. Altogether, these findings suggest that eccentric cycling is more taxing the brain than concentric cycling does.

A neuroimaging modality used in the study of Kan et al.^9^ was functional near-infrared spectroscopy (fNIRS). Functional NIRS is an optical neuroimaging technique based on the theory of neurovascular coupling^12^ and is relatively robust to motion artifacts.^13^ Since the local supply of oxygen is greater than its consumption in activated brain regions, a higher concentration of oxy-haemoglobin (O_2_Hb) and a decreased concentration of deoxy-haemoglobin (HHb) are to be observed.^13^ Changes in O_2_Hb and HHb recorded with 2 channels over the prefrontal cortex in Kan et al.^9^ were found to be smaller during two eccentric cycling exercises compared to a concentric cycling exercise, while there were higher scores for the two eccentric cycling exercise regarding the mental demand subscale of NASA-TLX. These preliminary findings suggest that a reduced supply of oxygen compared to relative mental demand occurs during eccentric cycling. However, they seem not to be supported by previous studies comparing the influence of the mode of muscle contractions from isolated and small muscle groups on brain activity.^10^ Using electroencephalography during elbow flexion at matched torque, a greater amplitude of movement-related cortical potentials in eccentric compared to concentric contractions was shown for electrodes located over the motor areas, prefrontal cortex and parietal lobe.^14^ Using functional magnetic resonance imagery (fMRI), Yao et al.^15^ observed an increased blood-oxygen-level dependent signal in similar brain areas during an eccentric task of the first dorsal interosseous muscle as compared to a concentric task. Yet, none of the neuroimaging studies mentioned above directly investigated cortical activity over multiple brain areas during whole-body eccentric and concentric exercise, such as cycling.

According to the aforementioned studies, the control of eccentric contraction would require important activation of the prefrontal regions, the right parietal lobe as well as the motor regions. These different areas are part of the central executive network known to be involved in cognitive-motor tasks modulating attentional demand^16^ as well as in the modulation of mental demand during continuous cycling exercise.^17^ As attention-demanding tasks exhibit increased activity in the prefrontal cortex and posterior parietal cortex, we hypothesized that the greater mental demand of eccentric cycling compared to concentric cycling would be associated with cortical activity increases in bilateral prefrontal cortices and right posterior parietal cortex. Finally, most of the aforementioned studies investigating eccentric cycling used an experimental design imposing an absolute mechanical power output. Using effort perception for matching the exercise intensity between eccentric and concentric cycling can be an alternative to circumvent the comparison of neurophysiological responses, either at the same mechanical or at the same metabolic workload.^18^ An effort perception approach to prescribe eccentric cycling has been proposed for the tailoring of rehabilitation programs in deconditioned populations, such as patients living with cardio-respiratory chronic diseases leading to muscle weakness.^3^ Considering the gaps in the literature, the main objective of the present study was to quantify the acute changes of attention-related cortical activity as well as those of relevant brain motor areas from fNIRS during submaximal eccentric and concentric cycling exercise. As the method to prescribe exercise is likely to influence cortical activity and mental demand, eccentric and concentric cycling was performed at both matched perceived effort and torque.

## METHODS

### Participants

Nineteen healthy male adults (mean age: 30 ± 6 years) volunteered in the study. All participants reported no sign of neurological, respiratory and cardiovascular disease or medication, which might affect brain and muscle functions. Participants provided written informed consent prior to participation in the study. The study was approved by the local ethics committee (ID: RCB_2016-A01722-49) and was conducted in accordance with the ethical standards specified by the Declaration of Helsinki. This study is belonging to the ExoMode project registered on ClinicalTrials.gov (NCT03280875). The sample size required for the present study was calculated using G*Power (3.1.9.6). The fNIRS results between the concentric and eccentric cycling conditions in one previous study^9^ were used as group parameters. The power analysis indicated that a total of at least 17 participants was required for the brain activation (f = .85 – np2= ∼.42 , corresponding to large effect – ; α = .05; 1-β = .90). Accordingly, a sample size of 17 participants was recruited. Two additional participants were included in case of experimental outliers.

### Experimental procedures

Participants were requested to visit the laboratory twice. The first visit consisted of a familiarization session and the second visit was the experimental session. Data collection occurred solely in the experimental session.

### Familiarisation session

The first visit to the laboratory was a familiarisation session with a double objective. First, participants were familiarised with the CR100® scale to evaluate their perception of effort and quadriceps pain, as well as to self-regulate the cycling exercise intensity based on a fixed perceived effort intensity. Second, the participants experimented with pedalling, driven by a motor either clockwise (concentric cycling) or anticlockwise (eccentric cycling), on an isokinetic semi-recumbent prototype cycle ergometer (Tech Med TMS, 89290 Champs-sur-Yonne, France). The seat of the semi-recumbent cycle ergometer allows for greater torso stabilization compared with a normal upright cycling position. This familiarisation cycling session consisted of participants understanding that they need to resist the pedals moving anticlockwise while they would have to accompany the pedals moving clockwise by exerting a suitable force to match a target line appearing on a screen in front of them. Participants were asked to perform eccentric and concentric cycling in a well-coordinated manner such that the knee extensors of each leg are activated at the right timing and intensity to ensure smooth pedal rotation and torque production. This familiarization session was carried out with the participant reaching submaximal torque values indicated by visual feedback on the control screen. The visual feedback was adjusted to produce 75W. To ensure precise execution of the concentric and eccentric isokinetic exercise, the researcher employed manual palpation of the quadriceps as a mean of verification. Specifically, this involved the researcher using manual palpation of the quadriceps to confirm that participants were adequately familiarized with the task and successfully contracted their quadriceps at the appropriate timing during pedal rotations. Finally, the participants performed four submaximal cycling exercises at a moderate intensity for three minutes at 30 RPM. The mode of muscle contraction (eccentric vs concentric) and the prescription of the exercise (fixed perceived effort or torque) were manipulated as follow: 1) eccentric and 2) concentric cycling at a moderate effort intensity (corresponding to 23 on the CR100® scale); 3) eccentric and 4) concentric cycling at a fixed torque value. The familiarization session was carried out at least 48 hours before the experimental session.

### Recording session

In this second session, participants performed four cycling exercise conditions for 5 min at 30 RPM (see Figure 1), as experienced in the familiarisation session. The choice of the exercise – rest ratio of 5 min – 5 min is based on pilot testing as well as the literature. First, as the literature demonstrated that 5 min of concentric or eccentric cycling is sufficient to obtain a steady state in physiological responses^4,5^, we decided to use this duration for the cycling bouts. Second, we tested this ratio during a pilot experiment in 5 participants to ensure that completion of our experimental protocol did not induce a marked decrease in maximal voluntary isometric contraction peak torque (MVIC) of the knee extensor (i.e., neuromuscular fatigue), a confound factor known to increase perception of effort.^19,20^ This pilot experiment consisted of duplicating the experimental session without measuring neurophysiological variables. Before the test session, MVIC of the dominant knee extensors was assessed (under verbal encouragement) as the best performance in three 5-s trials separated by a 30-s resting period. Then, one MVIC was performed in less than 20-s after the completion of the test. MVIC was performed on an isokinetic dynamometer (System Pro 4, Biodex Medical System, New York, NY) at a knee angle of 90°. We observed no marked decrease in knee extensor MVIC before starting the protocol (pre) vs after (post) completion of the protocol [MVC pre = 318 ± 77 Nm; MVC post = 318 ± 80 Nm; p = 0.982. d (95%CI) = −0.011 (−0.887, 0.867). Finally, to control for potential muscle damage at the onset of the experimental visit, we asked the participants at their arrival to the laboratory for the presence of muscle pain in their lower leg at rest or while walking, and no pain was reported.

**Figure 1.**
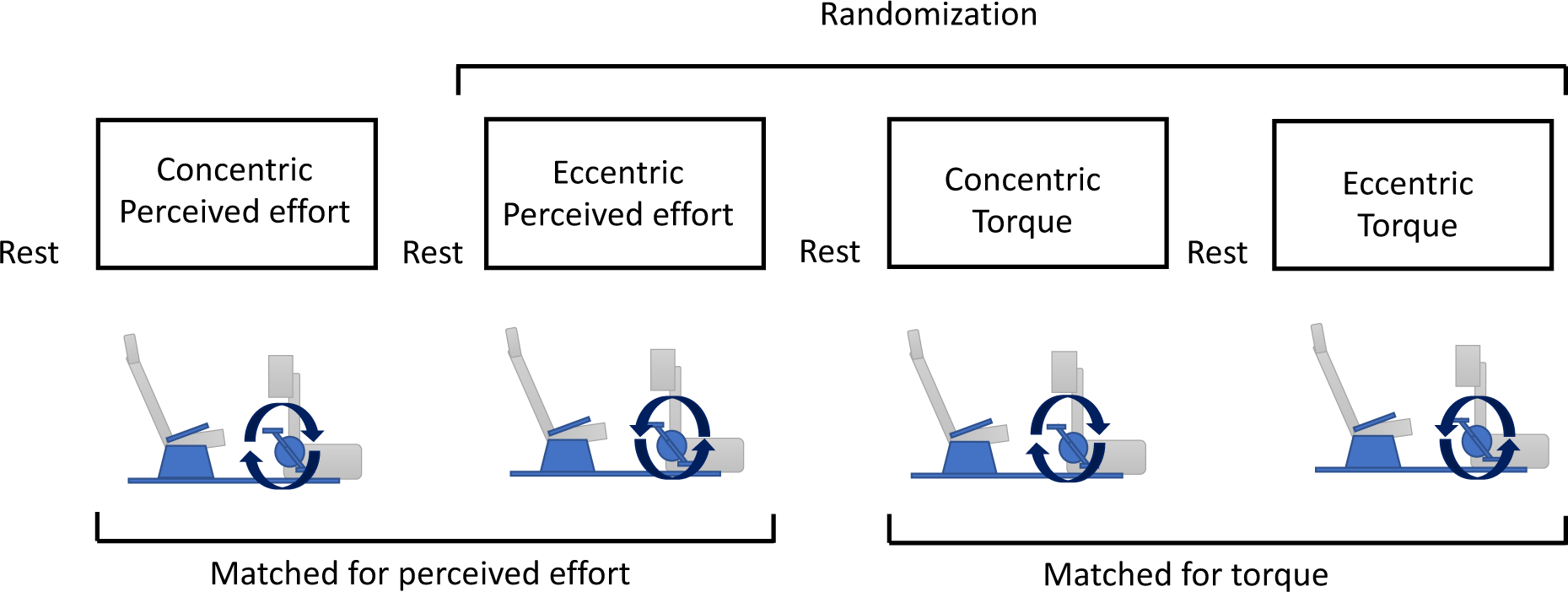
Overview of the experimental protocol. Four different 5-min cycling exercise conditions were realised in one session. In the first condition, participants performed concentric cycling at a fixed perception of effort (23 on the CR100® scale) without torque feedback. Then, the other three conditions were performed in a randomised order: eccentric cycling at a fixed perception of effort (23 on the CR100®), concentric or eccentric cycling at an intensity corresponding to the average torque produced in the first condition. Five minutes of rest were allowed between each exercise condition.

In the first condition, participants performed concentric cycling at a fixed perception of effort intensity corresponding to 23 on the CR100® scale (moderate intensity). No force feedback was provided to the participants who were instructed to self-regulate their force during each pedal revolution solely on their effort perception intensity, and to keep it moderate during the 5-min of exercise. The maximal torque values produced at each pedal revolution were extracted from the cycle ergometer every 15 s from minutes 2 to 5 of the exercise. These values were then averaged over the last 3 minutes, excluding the last 15 s (i.e., when the participants were reporting their perception), to individually determine the target torque used in the subsequent conditions where exercise intensity was prescribed with a fixed torque. The same data points were considered for analysis in all conditions. Then, the other three conditions were performed in a randomised order: (i) concentric cycling at an intensity corresponding to the average torque produced in the first condition, (ii) eccentric cycling at an intensity corresponding to the average torque produced in the first condition, (iii) eccentric cycling at a fixed perception of effort intensity corresponding to 23 on the CR100® scale (moderate, same perception of effort intensity than the first condition). Five minutes of rest were allowed between each exercise condition so that the physiological responses (heart rate and fNIRS signals) return to baseline levels. Heart rate was recorded with a chest belt and heart rate monitor (Polar V800, Finland) during rest and exercise periods for each condition. The average heart rate was obtained during the resting period before the onset of exercise and over the last minute of each exercise condition.

### Psychological variables

Perception of effort, also known as perceived exertion or sense of effort, can be described as a cognitive feeling of work associated with voluntary actions.^20^ Participants were asked to rate their perception of effort and quadriceps pain (CR100® scale) and affective valence during the last 20 seconds of each condition. At the beginning of the first sessions, participants were provided with validated instructions to dissociate between perceptions of effort and muscle pain^21^ and had the possibility to ask questions to the researchers if clarification needed. In-task affective valence (i.e., the pleasure/displeasure one feels) was measured using the Feeling scale, an 11-point bipolar scale ranging from −5 to +5. Seven anchors are provided ranging from “very good” (+5) to ‘‘very bad’’ (−5). Finally, participants were asked to complete at the end of each condition two subscales (mental demand and physical demand) of a commonly used subjective workload level questionnaire, the NASA-TLX scale. For each subscale, participants had to answer from 0 (very low or high) to 100 (very high or low). The NASA TLX scale was provided to the participants immediately after cessation of the 5-min cycling bouts. The NASA TLX scale was completed within 1 min post cycling bout for all participants.

### Cortical activity

#### Recording

A two-wavelength (765 and 856 nm) continuous-wave 15-channels fNIRS system (Oxymon MkIII, Artinis Medical Systems, The Netherlands) was utilised to measure the changes in O_2_Hb and HHb over regions of interest (ROI): left and right dorsolateral posterior parietal cortex, right parietal lobe and motor regions on both hemispheres as a whole. The O_2_Hb and HHb data were acquired during rest and exercise periods at a sampling rate of 10 Hz. O_2_Hb and HHb concentration changes (expressed in ΔμM) obtained by using the modified Beer-Lambert law were displayed on a PC monitor in real time. The optodes (5 receivers and 13 transmitters resulting in a total of 15 channels, Figure 2) were placed on the participants using a fit neoprene fNIRS head cap to get a stable optical contact with the scalp by generating an equal amount of pressure. The pressure was adequate to induce a partial transient blockage of the skin circulation during the fNIRS study^22^ as witnessed by the presence of well-defined circles over the skin that disappears about 15 min after the test session. In the 15 channels, the measurement points were defined as the midpoint of the corresponding illuminator-detector pairs interspaced by 3 cm. The Brodmann’s areas covered by the different channels were extracted via the NFRI function^23^ from the MNI (Montreal Neurological Institute) coordinates (Figure 2). Cartesian coordinates were obtained with a Fastrak® 3D Digitizer (Polhemus Inc., USA) allowing the localization of each optode and channel from four anatomical reference points: nasion, right and left mastoids and vertex.

**Figure 2.**
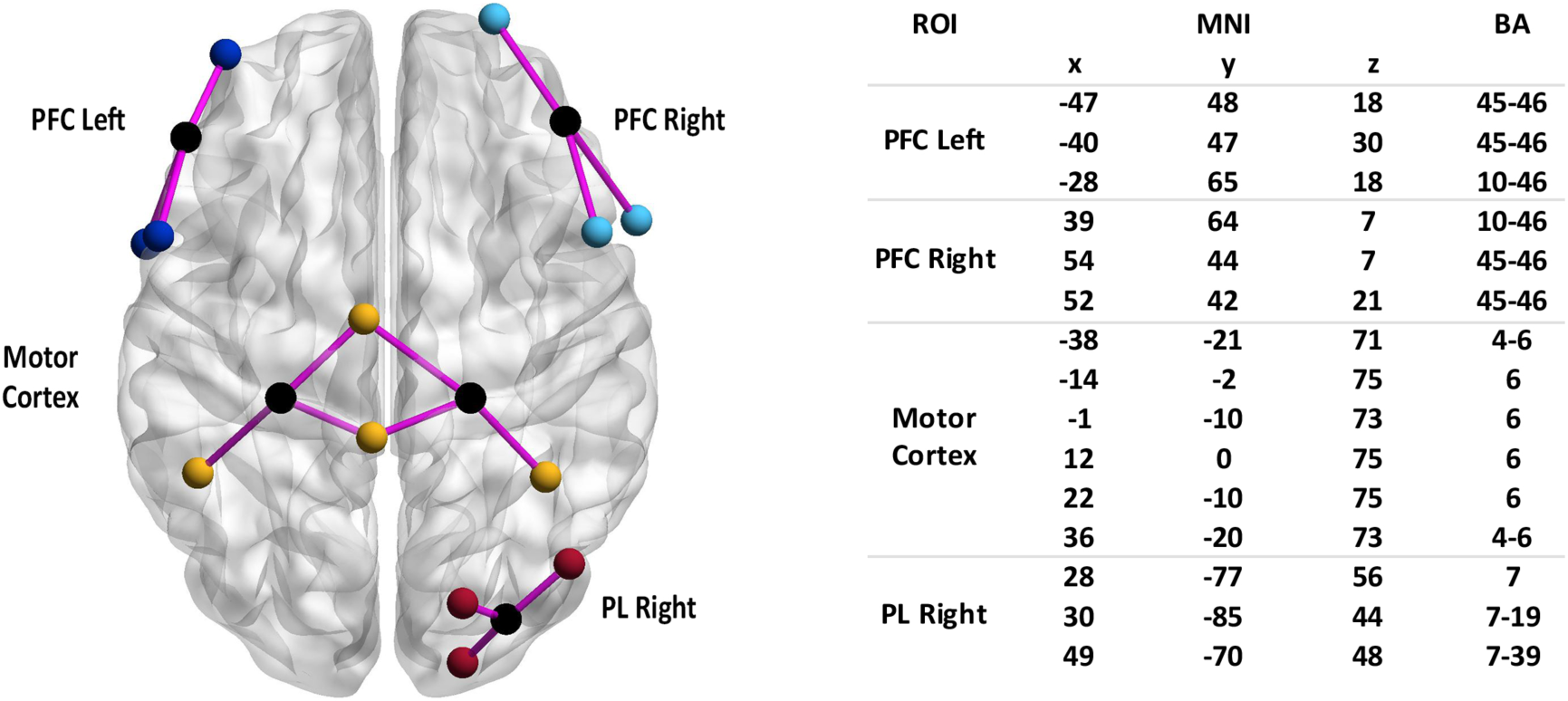
The functional near infrared spectroscopy (fNIRS) multichannel layout with 15 channels covering four regions of interest (ROI): motor cortex (yellow), left (dark blue) and right (light blue) prefrontal cortex (PFC) and parietal lobe (PL red). On the left, spatial organization of different optodes (BrainNet viewer toolbox^24^); in black the receivers, in colour the emitters, and pink edge the channels. On the right, MNI coordinates to each fNIRS channel (ROI) with corresponding Brodmann areas (BA).

#### Data analysis

Raw data (optical intensity) extracted from Oxysoft software (v3.0.75, Artinis Medical Systems) was converted in optical density (OD) using Matlab® (MATLAB 2014b, The MathWorks, USA). Then, to reduce possible head motion artifacts two methods implemented in the Homer2 toolbox^25^ were used. First, the moving standard deviation and spline interpolation methods were applied using parameters: SDTresh = 20, AMPTresh = 0.5, tMotion = 0.5s, tMask = 2s and p = 0.99. Second, we used wavelet-based signal decomposition with the parameter: iqr = 0.1, as recommended.^9^ Then OD time series were converted into relative concentrations of O_2_Hb and HHb using the modified Beer-Lambert law corrected by a differential pathlength factor depending on the age of the participant.^26^ A low-pass Butterworth zero-phase digital filter (order 4) with a cut-off frequency of 0.08 Hz was applied to limit physiological artifacts (heart rate: ± 1 Hz, respiration: ± 0.4 Hz and Mayer wave: ± 0.1 Hz). For each participant, channels not having a sufficient signal-to-noise ratio were excluded from the analyses after observation of the power spectrum density of the O_2_Hb signals where the presence of a cardiac rhythm in the signal (peak around 1 Hz) indicates good contact between the optodes and the scalp.^27^ After this preprocessing step, 14% of the channels were excluded.

Following a splitting of the fNIRS signals from events indicating the beginning of each condition, the average between −30 seconds and −10 seconds of the minute of rest preceding each condition was subtracted from the subsequent data during the exercise period. The following dependent variables attesting to exercise-related cortical activation magnitude were calculated per channel for the period (representative of the steady-state level) between 2 minutes and 4 minutes 30 seconds of exercise for each condition: area under the curve of O_2_Hb and HHb and average values. The average of the channels positioned on 4 ROIs (the right and the left posterior parietal cortex and primary motor, and right parietal lobe) was extracted for each participant and each condition.

### Statistical analysis

Statistical analyses were performed using Statistica (7.1, Statsoft Inc., France). Data were examined for normality and sphericity using Shapiro-Wilk and Mauchly tests, respectively. Where Mauchly’s tests indicated violations of the sphericity assumption, Greenhouse–Geisser corrections were applied. A one-way repeated-measure ANOVA was used to assess the differences between the four exercise conditions for heart rate recorded during the rest period. A fully repeated measures two-way ANOVA with the factors *mode of muscle contraction* (eccentric vs. concentric) and *mode of exercise prescription* (torque vs. perception of effort) were used to compare the NASA-TLX scores and fNIRS responses. In case of significant effect, post hoc comparisons were adjusted with the Bonferroni correction.

Wilcoxon signed-rank tests were performed for quadriceps pain and the Feeling scale values when comparing the mode of muscle contraction and the mode of exercise prescription. Student paired *t*-tests were carried out between the concentric and eccentric torque conditions for the perceived effort. For the cycling torque data, Wilcoxon signed-rank test was performed between the first condition (concentric at fixed perceived effort) and the two conditions (eccentric and concentric) where the intensity was fixed by a torque value.

Effect sizes were reported with the partial eta square (η^2^_p_) for the effects of the ANOVAs^28^ and with Cohen’s *d* and rank-biserial correlation (rrb) for pairwise comparison. The significance level for all analyses was established at *p* = .05 (two-tailed). All the fNIRS *p*-values were corrected using false discovery rate to control for type 1 error due to the spatial dependency between brain areas.

## RESULTS

### Torque and heart rate

The average torque during concentric cycling at a moderate perceived effort was 41.1 ± 14.1 Nm. No difference was observed between the average torque produced when performing concentric cycling at a moderate perceived effort, and the average torque produced when performing concentric (42.7 ± 13.8 Nm; W (17) = 47.5; p = .17; rrb = .33) or eccentric cycling at a fixed torque (40.3 ± 13.12 Nm; W (17) = 40; p = .08; rrb = .42).

No difference in resting heart rate was observed before the start of each cycling exercise condition (*F* (3, 45) = 1.25; *p* = .30; η^2^_p_ = .08). During the cycling exercises, the main effect of the mode of muscle contraction revealed a higher heart rate during concentric cycling compared to eccentric cycling (*F* (1, 18) = 40.3; *p* ˂ .001; η^2^_p_ = .69, 101 ± 13 vs. 87 ± 13 bpm). No main effect of the mode of exercise prescription (*F* (1, 18) = .09; *p* = .77; η^2^_p_ = .00; perception of effort: 94 ± 14 bpm; torque: 94 ± 15 bpm) or mode of muscle contraction × mode of exercise prescription interaction (*F* (1, 18) = .88; *p* = .36; η^2^_p_ = .05) was observed.

### Psychological variables

When the exercise was prescribed with a fixed torque feedback, the *perception of effort intensity* was higher (*t* (17) = 3.39; *p* ˂ .01; *d* = 1.07) during concentric cycling compared to eccentric cycling (23.2 ± 3.9 and 17.4 ± 6.8).

No main effects of the mode of muscle contraction (*W* (38) = 96.5; *p* = .75; r_rb_ = .05) and prescription of intensity (*W* (38) = 53; *p* = .44; r_rb_ = .13) were found for perceived quadriceps pain during the exercise as well as for the affects (Feeling scale) (*W* (38) = 94; *p* = .68; r_rb_ = .07 and *W* (38) = 79; *p* = .12; r_rb_ = .25, respectively).

For the perceived measures with the NASA-TLX subscales, one participant did not complete the questionnaire at the end of the experimental session. Therefore, data are available for 18 participants. For the *mental demand scores*, the main effect of the mode of muscle contraction revealed a higher mental demand during eccentric cycling compared to concentric cycling (*F* (1, 17) = 7.86, *p* = .01; η^2^_p_ = .32; Figure 3a). The main effect of the mode of exercise prescription revealed a higher mental demand when the exercise was prescribed with a fixed torque compared to a fixed perception effort (*F* (1, 17) = 26.22; *p* ˂ .001; η^2^_p_ = .61; Figure 3b). No mode of muscle contraction × mode of exercise prescription interaction was observed (*F* (1, 17) = .45; *p* = .51; η^2^_p_ = .02). For the *physical demand scores*, the main effect of the mode of muscle contraction revealed a higher physical demand during concentric cycling compared to eccentric cycling (*F* (1, 17) = 6.66; *p* = .02; η^2^_p_ = .28; Figure 3c). No main effect of the mode of exercise prescription was observed (*F* (1, 17) = .54; *p* = .47; η^2^_p_ = .03; observed power = .11). Follow-up tests on the mode of muscle contraction × mode of exercise prescription interaction (*F* (1, 17) = 6.99; *p* = .02, η^2^_p_ = .29) are presented in Figure 3d. The physical demand was greater for the torque condition in concentric cycling compared to eccentric cycling (*p* = .005, *d* = 0.315).

**Figure 3.**
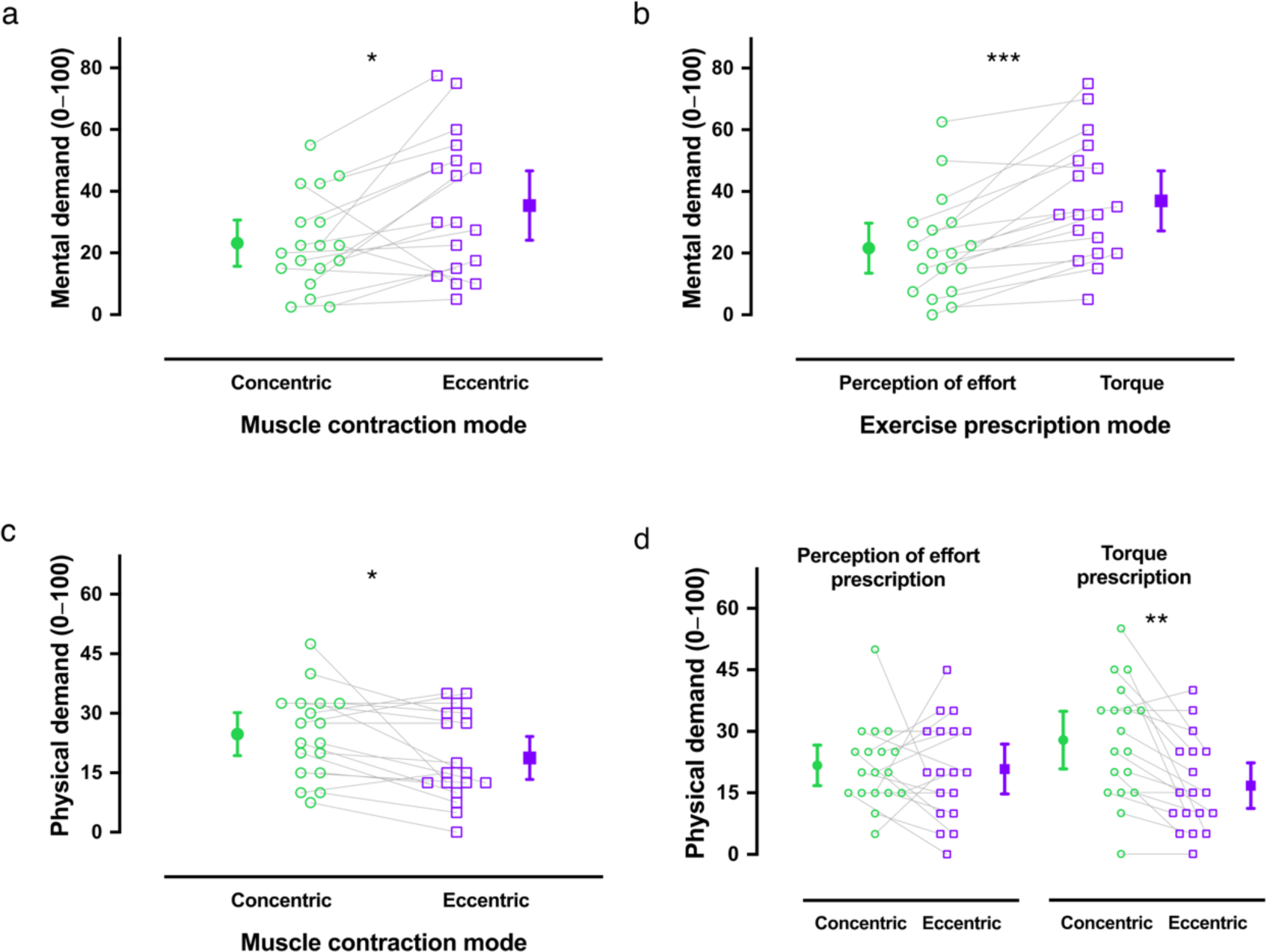
Perceived mental and physical demand scores measured with the NASA-TLX scale. The mental demand was perceived higher during eccentric cycling compared to concentric cycling (main effect of the muscle contraction mode, panel a), and during the exercise prescribed with the torque feedback compared to the perception of effort (main effect of the exercise prescription mode, panel b). The physical demand was perceived lower during eccentric cycling compared to concentric cycling (main effect the muscle contraction mode, panel c), and the interaction effect revealed a specific lower perceived physical demand during the eccentric cycling compared to concentric cycling when exercise was prescribed with the torque feedback. Concentric cycling and eccentric cycling are represented with circle and square markers, respectively. Individual (n=18) data are represented with empty markers and mean (±95% CI) as filled markers. * *p* < .05, ** *p* < .01, *** *p* < .001.

### Cortical responses

For the *left prefrontal cortex* (Figure 4A) and O_2_Hb, the main effect of the mode of muscle contraction revealed a higher area under the curve during eccentric cycling compared to concentric cycling (*F* (1, 18) = 16.35; *p* < .01, η^2^_p_ = .48). Neither the main effect of the mode of exercise prescription method (*F* (1, 18) = .1; *p* = .75; η^2^_p_ = .01), nor a mode of muscle contraction × mode of exercise prescription interaction (*F* (1, 18) = 1.48; *p* = .24; η^2^_p_ = .08) was observed. Regarding HHb, the main effect of the mode of muscle contraction revealed a lower area under the curve during eccentric cycling compared to concentric cycling (*F* (1, 18) = 35.77, *p* ˂ .001, η^2^_p_ = .67). Neither the main effect of the mode of exercise prescription (*F* (1, 18) = .49, *p* = .49; η^2^_p_ = .03) nor a mode of muscle contraction × mode of exercise prescription interaction (*F* (1, 18) = .07; *p* = .80; η^2^_p_ = .003) was observed.

**Figure 4.**
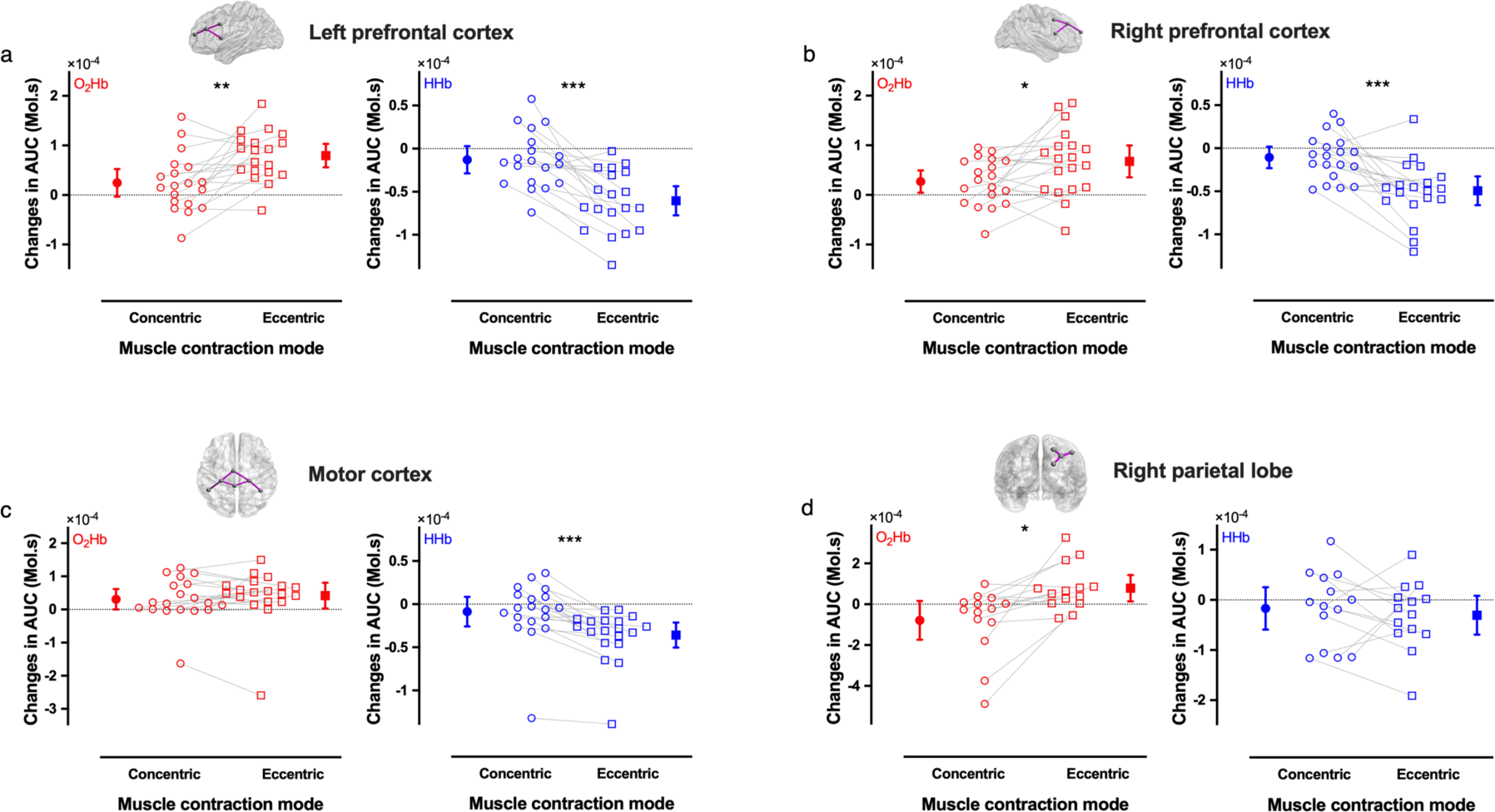
Changes in the area under the curve (AUC) for the oxyhaemoglobin (O2Hb, red) and deoxyhaemoglobin (HHb, blue). Data are presented for the main effect of the muscle contraction mode. The four regions of interest were the left prefrontal cortex (N=19, panel a), the right prefrontal cortex (N=19, panel b), the motor cortex (N=19, panel c) and the right parietal lobe (n=14, panel d). Data missing for the right parietal lobe is due to a damaged optical fiber occurring during one day of the data collection. Concentric cycling and eccentric cycling are represented with circle and square markers, respectively. Individual data are represented with empty markers and mean (±95% CI) as filled markers. * *p* < .05, ** *p* < .01, *** *p* < .001.

For the *right prefrontal cortex* (Figure 4B) and O_2_Hb, the main effect of the mode of muscle contraction revealed a higher area under the curve during eccentric cycling compared to concentric cycling *(F* (1, 18) = 6.96; *p* < .05; η^2^_p_ = .28). Neither the main effect of the mode of exercise prescription (*F* (1,18) = .30; *p* = .59; η^2^_p_ = .02) nor a mode of muscle contraction × mode of exercise prescription interaction (*F* (1, 18) = 2.06; *p* = .17; η^2^_p_ = .10) was observed. Regarding HHb, the main effect of the mode of muscle contraction revealed a lower area under the curve during eccentric cycling compared to concentric cycling (*F* (1, 18) = 22.26, *p* ˂ .001, η^2^_p_ = .55). Neither the main effect of the mode of exercise prescription (*F* (1,18) = .63, *p* = .44; η^2^_p_ = .03) nor a mode of muscle contraction × mode of exercise prescription interaction (*F* (1, 18) = 3.15, *p* = .09; η^2^_p_ = .15) was observed. For bilateral *motor cortex* and O_2_Hb (Figure 4C), no main effect of the mode of muscle contraction was observed (*F* (1,18) = 1.01, *p* = .33; η^2^_p_ = .05). Neither the main effect of the mode of exercise prescription (*F* (1,18) = .18, *p* = .68; η^2^_p_ = .01) nor a mode of muscle contraction × mode of exercise prescription interaction (*F* (1, 18) = .37, *p* = .55; η^2^_p_ = .02) was observed. Regarding HHb, the main effect of the mode of muscle contraction revealed a lower area under the curve during eccentric cycling compared to concentric cycling (*F* (1, 18) = 38.26, *p* ˂ .001, η^2^_p_ = .68). Neither the main effect of the mode of exercise prescription (*F* (1, 18) = .73, *p* = .40; η^2^_p_ = .04) nor a mode of muscle contraction × mode of exercise prescription interaction (*F* = (1, 18) = 3.42, *p* = .08; η^2^_p_ = .16) was observed.

For the *right parietal lobe* (Figure 4D) and O_2_Hb, the main effect of the mode of muscle contraction revealed a higher area under the curve during eccentric cycling compared to concentric cycling (*F* (1, 13) = 11.12, *p* = .02, η^2^_p_ = .46). Neither the main effect of the mode of exercise prescription method (*F* (1,13) = 1.09, *p* = .32; η^2^_p_ = .08; Figure 4D) nor a mode of muscle contraction × mode of exercise prescription interaction (*F* (1, 13) = .20, *p* = .66; η^2^_p_ = .01) was observed. Regarding HHb, no main effect of the mode of muscle contraction was observed (*F* (1, 13) = .38, *p* = .55; η^2^_p_ = .03). Neither the main effect of the mode of exercise prescription (*F* (1, 13) = 2.2, *p* = .16; η^2^_p_ = .14 nor a mode of muscle contraction × mode of exercise prescription interaction (*F* (1, 13) = 1.33, *p* = .27; η^2^_p_ = .09) was observed.

## DISCUSSION

The main objective of the present study was to quantify the acute changes in attention-related cortical activity as well as those relevant to brain motor areas from fNIRS during eccentric and concentric cycling exercises performed at both matched perceived effort (23 on CR100® scale) and torque. The main results showed a greater mental demand and cortical activation in the right and left prefrontal cortices and right parietal lobe during eccentric cycling compared to concentric cycling. These findings confirm previous results suggesting that eccentric cycling is more cognitively demanding than concentric cycling^9^ and extend them by identifying specific brain activation patterns underlying this greater mental demand.

### Mental demand during eccentric cycling

Mental demand was subjectively measured with the NASA-TLX subscale. The mental demand was reported to be 45% greater during eccentric cycling compared to concentric cycling, while physical demand was 38% greater during concentric cycling. These findings support the results of Kan et al.^9^ Nevertheless, mental and physical demand values were 2 to 3 times lower in the present study. The higher mental and physical demand found in Kan et al.^9^ is likely due to a greater exercise intensity compared to the present study, as confirmed by the higher mean heart rate values reported in their study as compared to ours (eccentric cycling: ∼+29 bpm; concentric cycling: +56 bpm). While mental demand refers to the mental and perceptual activity imposed by a task and refers to related information processing, effort refers to the voluntary engagement of mental and physical resources to perform, or attempt to perform, in a task. ^20,29^ In the present study, coordinating the leg movements properly during eccentric and concentric cycling required dedicated motor skills and so adequate demand for attention. As described in the methods, participants were asked to perform eccentric and concentric cycling in a well-coordinated manner to ensure smooth pedal rotation and torque production. This was similar to the cycling exercises proposed by Kan et al.^9^, where the authors observed that attention and vigilance were maintained after eccentric and concentric cycling. However, higher cognitive load (based on both the NASA TLX score and choice reaction time) was found in the study of Kan et al.^9^ during eccentric cycling. Altogether, it appears that eccentric cycling is more cognitively demanding than concentric cycling and that the less mental demand of eccentric and concentric cycling in the present study did not likely change cognitive functions post-exercise.

Regarding the mode of exercise prescription, the mental demand was significantly higher when the intensity was set by a target torque value compared to an exercise prescribed by a fixed perceived effort. To the best of our knowledge, this study is the first to observe this difference in mental demand. A lower mental demand during self-regulated exercise based on effort perception is not surprising as, in this specific condition, participants did not have at each pedal revolution to visually focus on a torque target to reach. Contrary to the mental demand, the mode of exercise prescription did not alter the reported physical demand of the cycling exercise. This observation is in line with no significant difference in heart rate between the two modes of exercise prescription.

### Cortical activity during eccentric cycling

Neuroimaging studies using various methods and exercise paradigms reveal inconsistencies in the role of the primary motor cortex in the production of eccentric muscle contractions.^10^ Yao et al.^15^ during unilateral movements of the first dorsal interosseous muscle in an MRI scanner, as Fang et al.^14^ in an EEG study for the elbow flexors, showed a greater activation of the primary motor cortex contralateral to the active muscles when performing eccentric compared to concentric contractions. Conversely, a reduction was reported in the primary motor cortex blood-oxygen-level dependent signal amplitude during eccentric compared to concentric contractions.^30^ The present study is the first to collect, during an eccentric cycling exercise (i.e., a bilateral task), fNIRS data over the precentral gyrus (BA 4/6) and primary motor areas (BA 4). The present study did not observe significant differences in motor cortices activity between eccentric and concentric cycling. Our results can be explained because the tasks performed are fundamentally different from the literature. First, the different studies cited above performed unilateral distal movements of the upper limb, unlike our study where the participants performed a bilateral task mobilizing larger muscle masses of the lower limbs. In addition, in the present study, the cycling exercise was performed at moderate intensity (23 on the CR100® scale or using an associated torque). Studies showing a greater activity of the contralateral primary motor during eccentric muscle contractions involved exercises performed either at maximum intensity^14^ or sub-maximum intensity fixed relative to maximal voluntary contraction force or body weight.^14,15^ Noteworthy, in the study of Olsson et al.^30^ a continuous imagery task paradigm was used by alternating concentric (lifting a weight) and eccentric (slowing down a weight) muscle contractions. For other cortical areas involved in the motor network (e.g. primary and secondary motor areas not recorded here), a consensus seems to exist on the greater cortical activity observed during eccentric contractions.^10^

The main results of the present study revealed a greater activity of the left and right prefrontal cortices, as well as the right parietal lobe, when performing eccentric cycling compared to concentric cycling, regardless of the mode of exercise prescription. These observations likely unveil underlying neurophysiological mechanisms sustaining the increased mental demand reported by the participants during eccentric cycling. The left and right prefrontal cortices as well as the right parietal lobe are known for their roles in the modulation of mental demand during prolonged exercise^17^ as well as during sustained attentional cognitive tasks.^16^ When focusing on a specific movement to perform, several brain areas, constituting the central executive network, are engaged. More specifically, these regions would be part of the fronto-parietal network that consists mainly of the dorsolateral prefrontal cortex, the frontal cortex and the parietal lobe.^31^ An increase in eccentric force variability has been proposed as a sign of an increased motor control difficulty during eccentric contractions compared to concentric contractions.^11^ Controlling the variability in motor output (not assessed in the present study) during eccentric contractions would likely require greater attentional demand, thus explaining the observed increase in the activity of the prefrontal cortices and the right parietal lobe. Some authors have highlighted the complementarity of regional brain activity magnitude measures and those linked to the dynamic activity of brain networks.^32^ On this basis, the fMRI study of Yao et al.^33^ examined the strength of functional connectivity between the primary motor cortex and other cortical regions in the motor control network (e.g. supplementary motor area, cerebellum) involved in eccentric and concentric contractions. The authors observed a weaker functional connectivity during eccentric compared to concentric contractions of the right first dorsal interosseous muscle. A weaker statistical dependence between two-time series from different brain areas may reflect more involvement of specialised regions (segregation) and thus less functional integration.^34^ This means that higher cortical activation may be required during eccentric contraction to compensate for a weaker functional connectivity. Yao et al.^33^ hypothesized that the brain control of a concentric contraction will be more efficient than an eccentric contraction. This first result needs to be supported by more evidence, notably for a bilateral task mobilizing larger muscle masses of the lower limbs. The analysis of functional connectivity could be a relevant and complementary approach to better understand the functional organisation of the brain when performing eccentric cycling over time and its possible implications for rehabilitation and training.

### Methodological considerations

In the present study, we controlled for the induction of muscle fatigue by providing a sufficient recovery period between cycling exercise conditions, thus allowing us to ensure that our results are not confounded by a decrease in force production capacity of the working muscles with changes in the corticospinal excitability.^35^ Exercise-induced muscle fatigue plays an important role in the modulation of cortical activity during exercise. In an fMRI study involving a maximal-effort handgrip contraction, Liu et al.^36^ observed an early increase followed by a decrease in brain activity in the primary motor cortex and in particular in the prefrontal cortex. The exercise intensity in our experimental design was low as indicated by perceived effort and heart rate values. In addition, our control experiment showed no significant decrease in the maximal force production capacity of the knee extensors after the whole exercise protocol.

The present study faces one limitation, which might be taken into account in follow-up studies. The fNIRS signal can be contaminated by the physiological responses to the task performed (e.g. heart rate, blood pressure). This signal contamination related to non-neural activity can lead to false positives or negatives in our analyses.^37^ To overcome these issues, the use of short-distance channels (approximately 1 cm between the transmitters and the receivers) was proposed.^38^ In this study, we did not use such short-distance channels for separating signals that stem from blood flow changes in the extracerebral layers (scalp blood flow) as well as the cerebral tissue layer (systemic blood flow). As indicated in the methods, we minimized this potential effect by inducing a partial transient blockade of the skin blood flow during the fNIRS data collection.^22^ Interestingly, heart rate was ∼13% higher in concentric as reported before^39^, whereas the greatest cortical activation was observed during eccentric cycling. These elements allow us to think that the results might not be influenced by systemic interference but future investigations are necessary to confirm this hypothesis.

Finally, it is important to note that 2-3 participants did not report a higher mental demand during eccentric cycling compared to concentric cycling. These interindividual variability responses may be due to the differences in the degree of expertise of our participants with eccentric cycling. Therefore, future studies should explore the link between the repeated and chronic use of eccentric cycling and a potential decrease in its mental demand. Regarding the variability in fNIRS responses, 3-5 participants presented results contradictory to the group. In line with this observed variability, we strongly encourage future researchers to increase their sample size beyond the one used in our present study to replicate and confirm our observations.

## CONCLUSION

In the present study we took advantage of using functional near infrared spectroscopy covering four regions of interest to investigate how mental demand induced by eccentric exercise modulated the central executive network. The present study showed that the higher mental demand observed during eccentric cycling compared to concentric cycling was associated with a greater activation of the left and right prefrontal cortices as well as the right parietal lobe. These findings provide evidence that heightened activation of the frontoparietal network may support eccentric movement control.

## PERSPECTIVES

A large proportion of our daily living activities require accurate control of movements employing eccentric contractions (e.g. walking downstairs, sitting). We encourage future studies to replicate and extend our results using different locomotion modalities such as uphill or downhill running, which are known to predominantly involve concentric and eccentric contractions of the knee extensors^40^, respectively. As physiological responses to physical exercise are well known to be linked to the exercise intensity, it could also be of interest to explore the interaction between the exercise intensity and mental demand during various exercise modalities. Based on the findings of this study, future studies should investigate the potential use of wearable fNIRS for clinical and sports applications via an online neurofeedback approach to modulate the mental demand. Further investigations are also warranted to examine the impact of task familiarization on mental demand and the related brain activation during eccentric cycling. As learning is associated with increased automaticity and decreased resources engaged in a task, it is plausible that participants will experience lower mental demand when they become acquainted with the task. Another application is that self-regulated eccentric exercise based on perceived effort is less mentally demanding, suggesting that it may be more appropriate if one session needs to minimize the overall workload. Overall, eccentric cycling workload can be prescribed based on the individual’s perceived rate of exertion, but also at a brain activation workload similar to that elicited by concentric cycling to improve both neuro-motor processing strategies and mental demand.

## ACKNOWLEDGEMENT

This research work was supported by the French National Research Agency (EXOMODE–ANR–15–CE19-0023). We thank the anonymous reviewers for their careful reading of our manuscript and their many insightful comments and suggestions.

## AUTHOR CONTRIBUTIONS

Lenaic Borot: Conceptualization; data curation; formal analysis; investigation; methodology; validation; visualization; writing –original draft; writing –review and editing. Benjamin Pageaux: Conceptualization; investigation; methodology; visualization; validation; writing –review and editing. Davy Laroche: Conceptualization; funding acquisition; methodology; resources; supervision; writing –review and editing. Gregoire Vergotte: methodology; formal analysis; writing –review and editing. Romuald Lepers and Stephane Perrey: Conceptualization; investigation; funding acquisition; methodology; project administration; resources; supervision; validation; writing – review and editing.

